# Structural specialities, curiosities and record-breaking features of crustacean reproduction

**DOI:** 10.1101/034223

**Authors:** Günter Vogt

**Affiliations:** Faculty of Biosciences, University of Heidelberg, Im Neuenheimer Feld 230, 69120 Heidelberg, Germany

**Keywords:** Crustacea, reproduction, structural peculiarities, curiosities, records

## Abstract

Crustaceans are a morphologically, physiologically and ecologically highly diverse animal group and correspondingly diverse are their reproductive characteristics. They have evolved structural specialities with respect to penis construction, sperm form, sperm storage, fertilization and brood care. Unique in the animal kingdom are safety lines that safeguard hatching and first molting. Further curiosities are dwarf males in parasitic and sessile taxa and bacteria-induced gigantism and infectious feminization in crustacean hosts. Record-breaking features in animals are relative penis length, clutch size, sperm size, chromosome number, viability of resting eggs, and fossil ages of penis, sperm and brooded embryos. These reproductive peculiarities are reviewed and their implication for basic and applied biology is discussed, including the early evolution and diversification of brood care in arthropods, sperm competition and assurance of paternity, posthumous paternity and sustainable male-based fishery, and ecotype changes by man-made pollution.

## INTRODUCTION

The Crustacea are renowned for their exceptional diversity in morphology, physiology, life history and ecology. Dating back to the Early Cambrian they had ample time for experimentation with form and function (Sepkoski, 2000; Harvey et al., 2012). The crustaceans include six classes, 42 orders, 849 extant families and approximately 60.000 described species (Martin and Davis, 2001). Maximum body length of the adults ranges from 0.1 mm in a tantulocarid ectoparasite to more than 1 m (including chelae) in the American lobster. Crustaceans occur in marine, limnic and terrestrial habitats from the polar region to the tropics and from lowlands to high mountains. Most crustaceans are free living but some groups are sessile or parasitic. Because of this enormous diversity there are a lot of special adaptations in this animal group.

The most curious adaptations in animals have probably evolved in context with reproduction, particularly with respect to behaviour. Examples are the sexually cannibalistic praying mantis females, which sometimes eat the males during mating (Lawrence, 1992), and the megapodid birds, which incubate their eggs by heat generated from microbial decomposition or volcanism (Dekker and Brom, 1992). However, there are also many morphological reproductive curiosities such as the antlers of deer (Price et al., 2005) or the love darts of gastropods (Chase and Blanchard, 2006).

The present article highlights structural specialities, curiosities and record-breaking features of reproduction in crustaceans and emphasizes implications for broader zoological issues. The topics discussed include different construction principles of copulatory organs, extraordinary sperm types, paternity assurance and posthumous paternity, safeguarding of hatching and first molting, brooding of posthatching stages, manipulation of sex and body size by parasites, and animal records in relative penis length, chromosome number, viability of resting eggs and fossil age of reproductive structures.

## HYDRAULIC VERSUS FORM-INVARIANT COPULATORY ORGANS

Most crustaceans transfer sperm by copulatory organs. This evolutionary legacy has led to some curiosities in sessile and parasitic groups as explained further down. In this chapter, I want to compare two different construction principles of copulatory organs, the extensible hydraulic penis in sessile Cirripedia and the form-invariant and permanently stiff copulatory organs in Decapoda.

The thoracican Cirripedia are unique among sessile animals as they use a penis for sperm transfer (Fig. 1A–D) (Barnes, 1992). They are hermaphrodites but usually do not self-fertilize. Their unpaired penis is greatly extensible and anatomically well suited for sweeping movements to search for functional females in the surroundings (Fig. 1B). It is extended and retreated by modulation of the turgor pressure in the longitudinally running inflatable haemolymph channels (Fig. 1D) (Klepal et al., 1972). The cuticle of the penis is thin and annulated (Fig. 1C, D) and combines mechanical strength with flexibility, facilitating length variation (Klepal et al., 2010). Searching movements are enabled by a compact layer of longitudinal musculature underneath the epidermis (Fig. 1D) (Klepal et al., 1972). Circular musculature is only found around the central ductus ejaculatorius, which serves for sperm ejection. The anterior portion of the penis is studded with rows of sensory setae (Fig. 1C), which are assumed to help in finding of sexual partners. Cirripeds are the animal record holders in relative penis length and stretchability. For example, the penis of *Cryptophialus minutus* is eight times longer than its body size (Neufeld and Palmer, 2008). The resting penis of *Semibalanus balanoides* is 8–13 mm long, depending on size of the animal, but can be stretched to about four times this length (Klepal et al., 1972).

**Fig. 1.**
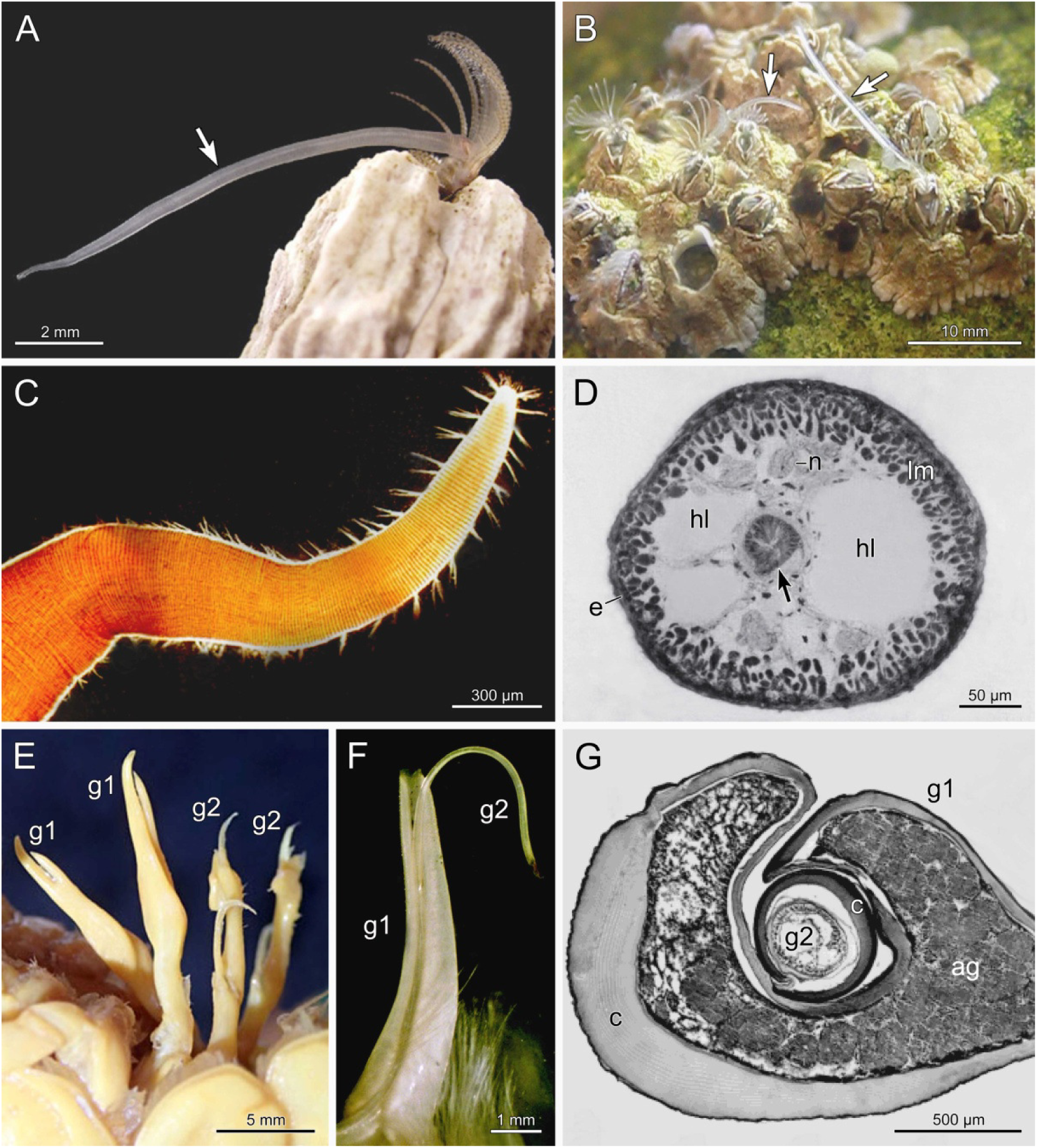
Hydraulic penis in Cirripedia (A-D) and form-invariant copulatory organs in Decapoda (E-G). (A) Relaxed penis (arrow) in *Balanus glandula* (photo: Christopher J. Neufeld). (B) Extended penises (arrows) in mating *Semibalanus balanoides* (from a video by Stefan Siebert). (C) Apical part of penis of *Semibalanus balanoides* showing annulations and groups of sensory setae (photo: J. Matthew Horch). (D) Cross section through penis of *Balanus balanus* showing inflatable haemolymph lacunae (hl). Arrow, ductus ejaculatorius; e, epidermis with thin cuticle; lm, longitudinal musculature; n, nerve (from Klepal et al., 1972). (E) Paired copulatory organs of crayfish *Orconectes cristavarius* consisting of first (g1) and second (g2) gonopods (photo: Tiffany Penland). (F) Copulatory organ of box crab *Calappula saussurei* in functional state with second gonopod inserted in first gonopod (from Ewers-Saucedo et al., 2015). (G) Cross section through copulatory organ of freshwater crab *Potamon gedrosianum* in functional state showing thick stabilizing cuticles (c) in both gonopods. ag, accessory sexual gland (from Brandis et al., 1999).

Most intertidal barnacles have a brief reproductive season and develop a functional penis only during this period (Barnes, 1992; Klepal et al., 2010). For example, in New York area, in which *Semibalanus balanoides* reproduces in late October and early November, the penis grows rapidly over September and October and degenerates again during November (Horch, 2009). Interestingly, intertidal barnacles can adapt the size and shape of the penis to local hydrodynamic conditions, as shown for *Balanus glandula* and *Semibalanus balanoides* (Neufeld and Palmer 2008; Horch, 2009). On wave-exposed shores, they develop shorter penises with greater diameters than in wave-protected sites and invest more energy and resources in penis development and function. Transplant experiments between wave-exposed and protected sites revealed phenotypic plasticity as the cause of penis variation rather than differential settlement or selective mortality, identifying environmental cues as determinants of genital diversification in animals aside of female choice, sexual conflict and male-male competition (Neufeld and Palmer 2008; Horch, 2009).

A totally different construction principle of the copulatory organs is found in the Decapoda. Most of these vagile and big-sized crustaceans possess permanent and form-invariant copulatory organs (Factor, 1995; Holdich, 2002; Becker et al., 2012). In freshwater crayfish and brachyuran crabs the paired copulatory organs are composed of the first and second pleopods (gonopods) (Fig. 1E, F), which serve different functions. The first gonopod forms a tube-like structure (Fig. 1G) that takes the sperm from the genital opening. The second gonopod is inserted into the tube of the first gonopod (Fig. 1F, G) and acts like a plunger, producing spermatophores of appropriate size for transfer to the female sperm storage site (Holdich, 2002; Becker et al., 2012). The form constancy and permanent stiffness of both gonopods is achieved by particularly thickened cuticles (Fig. 1G) (Brandis et al., 1999; Ewers-Saucedo et al., 2015). The complex structure and form-invariance qualifies the gonopods for taxonomic purposes. For example, in cambarid crayfish the morphology of the gonopods is the main criterion for species identification (Hobbs, 1989).

## GIANT SPERM AND EXPLOSION SPERM

The crustaceans have evolved an enormous variety of sperm forms and fertilization mechanisms (Jamieson, 1991). Some groups show the classical sperm morphology consisting of head and flagellum but most groups have derived sperms. Here, I want to discuss the two most curious types, the giant sperm of Ostracoda and the explosion sperm of Decapoda.

Ostracods of the superfamily Cypridoidea have some of the longest sperm in the animal kingdom (Fig. 2A–D), surpassed only by a few insects. Sperm lengths range from 268 μm to 11.8 mm corresponding up to 4.3 times the shell length of the producing male (Smith et al., 2016). The record holder in the animal kingdom is the fly *Drosophila bifurca*, which produces sperm of 58.2 mm, which is 20 times longer than the specimens that manufacture them (Pitnick et al., 1995). However, their length is mainly accounted for by the exceptionally long flagellum, whereas the aflagellate ostracod sperm is composed in its entire length by the nucleus and two enormous mitochondria. Although tailless, these spermatozoa are motile because contractile elements can produce longitudinal rotation of the entire sperm (Matzke-Karazs et al., 2014). Interestingly, despite their enormous length they enter the egg completely during fertilization (Matzke-Karasz, 2005).

**Fig. 2.**
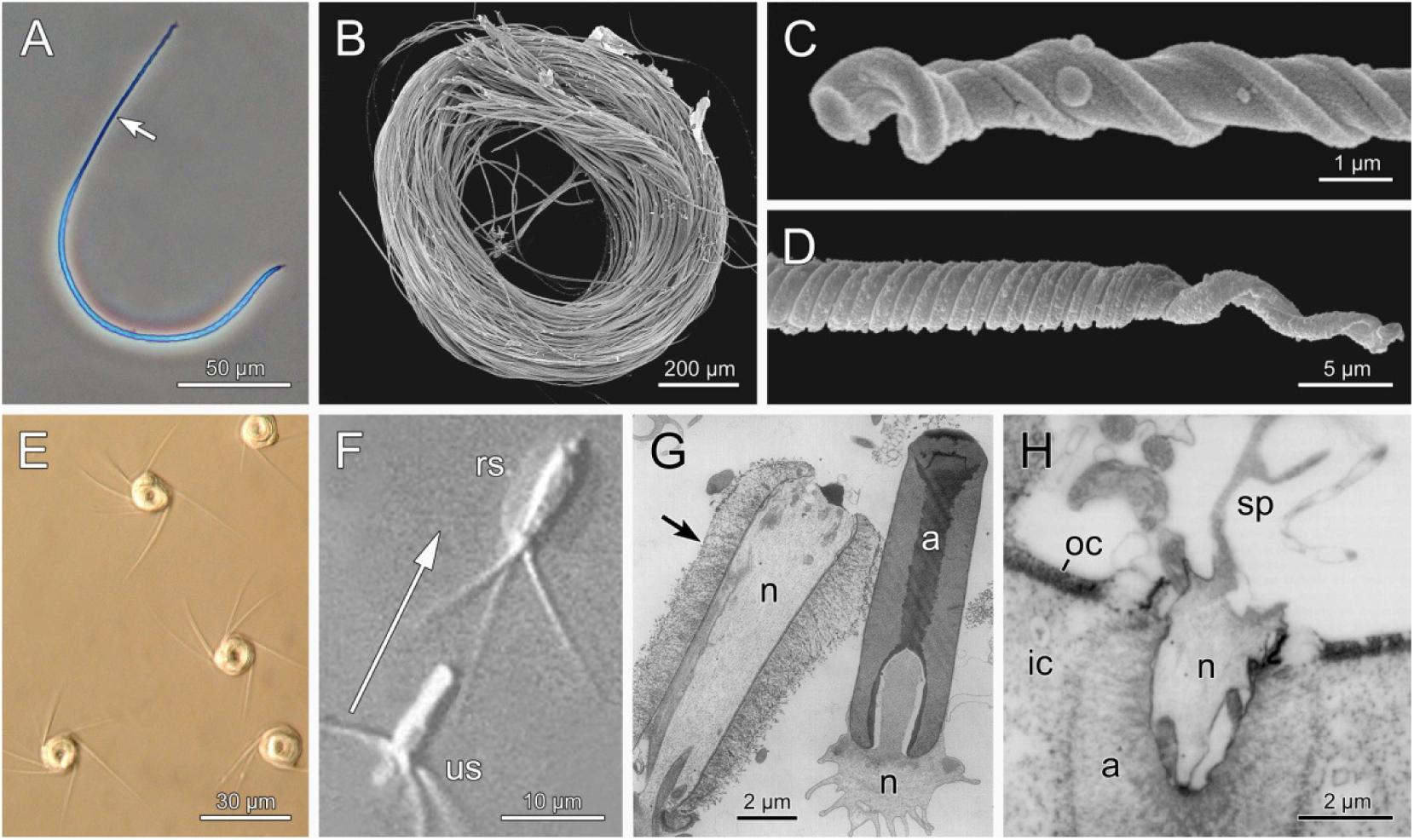
Giant sperm in Ostracoda (A-D) and explosion sperm in Decapoda (E-H). (A) Shortest giant spermatozoon from *Fabaeformiscandona velifera*. Arrow denotes anterior region (from Smith et al., 2016). (B) Longest giant spermatozoa in sperm bundle from seminal vesicle of *Australocypris robusta* (from Smith et al., 2016). (C) Drilled anterior tip of spermatozoon of *Pseudocandona marchica* (from Smith et al., 2016). (D) Coiled end piece of spermatozoon of *Eucypris virens* (from Smith et al., 2016). (E) Sperm of crayfish *Austropotamobius italicus* showing central body with nucleus and acrosome and extending radial arms (from Galeotti et al., 2012). (F) Movement (arrow) of spermatozoon of lobster *Homarus americanus* by abrupt acrosome eversion. Montage of two pictures from a video. rs, reacted sperm; us, unreacted sperm (from Tsai and Talbot, 1993). (G) Ultrastructural aspects of unreacted (right) and reacted (left) sperm of *Homarus americanus*. Note relative position of the nucleus (n) within the sperm. Arrow denotes eversed acrosomal material. a, unreacted acrosome (from Talbot and Chanmanon, 1980b). (H) Fertilization of oocyte. The acrosome is in the process of eversion and the nucleus is thereby torn through the envelope of the oocyte. ic, inner chorion layer; oc, outer chorion layer; sp, spike of spermatozoon (from Talbot et al., 1991).

The spermatozoa of reptantian Decapoda, which include freshwater crayfish, lobsters and crabs, are called "Explosionsspermien" (explosion sperm) in the German literature. Like the sperm of other decapods they are aflagellate and non-motile (Jamieson, 1991; Tudge and Koenemann, 2009). Reptantian sperm are composed of a compact central body consisting of the nucleus and acrosome, and radial arms (Fig. 2E) (Brown, 1966; Talbot and Chanmanon, 1980a; López-Camps et al., 1981; Vogt, 2002). The radial arms are rich in microtubules and become free when the spermatozoa are released from the spermatophore (Niksirat et al., 2014). Despite of the absence of flagellae, reptantian sperm can achieve short-term motility by an abrupt eversion of the acrosome. This reaction tears the nucleus from posterior to anterior and propels the entire sperm forward (Fig. 2F, G). In the American lobster *Homarus americanus* the explosive forward movement is about 18 μm (Fig. 2F) (Talbot and Chanmanon, 1980b). The force generated by acrosome eversion is sufficient to push the sperm nucleus through the egg coat (Fig. 2H) (Brown, 1966; Goudeau, 1982; Talbot et al., 1991).

## ASSURANCE OF PATERNITY BY SPERM DISPLACEMENT

Promiscuity of females and multiple paternity of clutches seems to be widespread in Crustacea. It is mainly known for Decapoda but there are also examples for Isopoda, Cirripedia and Copepoda (Avise et al., 2011; Dennenmoser and Thiel, 2015). Avise et al. (2011) compiled data on paternity in 11 decapod species including shrimps, crayfish, lobsters and crabs. In nine species, microsatellite analysis revealed multiple paternity in 20-100% of clutches. The mean number of fathers per clutch and species ranged from 1 to 5.3. The maximum number was 11 in the shrimp *Caridina ensifera* (Yue and Chang, 2010). In crayfish clutches fertilized by more than two males, one sire always dominated the brood by sharing his genes with 50-80% of the offspring (Walker et al., 2002; Yue et al., 2010; Kahrl et al., 2014). The pre-condition for multiple paternity is promiscuity of the female and storage of the sperm of all mates until egg-laying, which often occurs weeks, months or even years after the last mating (Jensen and Bentzen, 2012). The males can enhance their chance to sire the offspring by removing the sperm of earlier mates or displacing it to an unfavourable place.

Decapod females store sperm either externally as attached spermatophore clump (Fig. 3A) or internally in specialized seminal receptacles (Fig. 3D). Examples of the former are the astacid crayfish and examples for the latter are the eubrachyuran crabs. In the Astacidae, the spermatophores are glued to the sternal plate between the genital pores (Fig. 3A). Late males may leave the spermatophores of earlier males untouched or remove them partly or completely, which significantly influences paternity of the clutch. In laboratory reared *Austropotamobius italicus*, 33% of the males completely removed the spermatophores of previous rivals, 63% left intact some of the rivals’ spermatophores (Fig. 3B, C) and 3% simply added their own sperm (Galeotti et al., 2007). In species with external sperm storage paternity may also incidentally be biased by mechanical abrasion of spermatophores during locomotion (Galeotti et al., 2007).

**Fig. 3.**
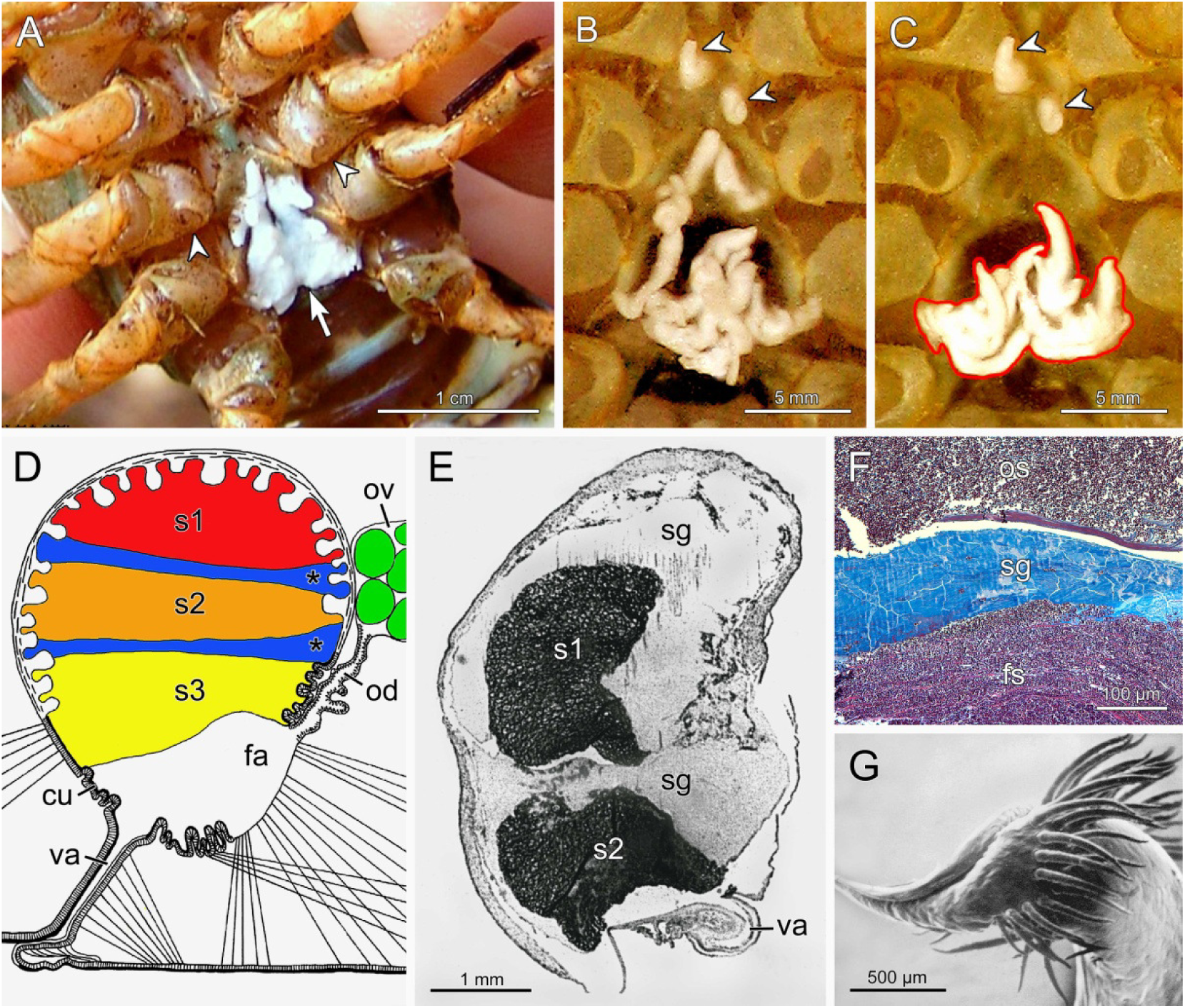
Storage and displacement of sperm in Decapoda. (A) Spermatophores attached to sternal plate (arrow) between female gonopores (arrowheads) in crayfish *Astacus astacus* (photo: Lukáš Konečný). (B, C) Partial removal of spermatophores of earlier mate (B) by following mate (C) in crayfish *Austropotamobius italicus*. Arrowheads denote persisting spermatophores and red line mark newly added sperm (from Galeotti et al., 2007). (D) Scheme of seminal receptacle of eubrachyuran crabs illustrating stratification of sperm packages (s1-s3) from three matings. The ejaculates are separated by wax-like sperm gel layers (asterisks). The sperm of the last male is deposited closest to the fertilization area (fa). cu, cuticle; od, oviduct; ov, ovary; va, vagina (based on a drawing by Becker et al., 2011). (E) Histological section through seminal receptacle of spider crab *Inachus phalangium* showing two sperm packages separated by sperm gel (sg) (from Diesel, 1990). (F) Older (os) and fresh (fs) sperm packages separated by sperm gel in cancrid crab *Metacarcinus edwardsii* (from Pardo et al., 2013). (G) Tip of first gonopod of snow crab *Chionoecetes opilio* with spoon-like structure and setal brushes suitable for the displacement of sperm from an earlier rival (from Beninger et al., 1991).

In eubrachyuran females, the sperm is stored in ventrally located paired seminal receptacles that are continuous with the oviduct and vagina (Fig. 3D) (Diesel 1991; Becker et al., 2011; McLay and López Greco, 2011; Pardo et al., 2013). The oocytes are internally fertilized in an area close to the opening of the oviduct (Fig. 3D). Since the spermatozoa of brachyurans are non-motile the sperm stored closest to the fertilization area have the highest chance to fertilize the eggs. Males may maximize their chance to sire the offspring by removing the sperm of predecessors from the seminal receptacle or by displacing it to an unfavourable position. The former variant is known from insects, which have specifically shaped gonopods with recurved spoon-like tips and swap-like bundles of setae (Waage, 1979; Tsuchiya and Hayashi, 2014). Beninger et al. (1991) found similarly structured gonopods in the snow crab *Chionoecetes opilio* (Fig. 3G) and stressed the possibility of sperm removal in this species. However, until now there is no experimental proof for active sperm removal in brachyurans.

In contrast, the displacement and sealing of the ejaculate of earlier mates is proven for several species (Diesel, 1990, 1991; Urbani et al., 1998; Sainte-Marie et al., 2000; Pardo et al., 2013). For example, in spider crab *Inachus phalangium* and cancrid crab *Metacarcinus edwardsii* mating males push the sperm packages of predecessors dorsally towards the apex of the receptacle and seal them off with a hardening wax-like sperm gel. They then place their own sperm closest to the oviduct opening (Fig. 3E, F) (Diesel 1990, 1991; Pardo et al., 2013). In snow crab *Chionoecetes opilio* up to 12 stratified ejaculate layers were found per seminal receptacle (Sainte-Marie et al., 2000). The wax-like material is probably produced in accessory sexual glands located in the basis of the first gonopods (Fig. 1G) (Beninger and Larocque, 1998; Brandis et al., 1999).

Mating experiments with the snow crab revealed that sperm from the most proximal sperm layer fertilizes all eggs of a clutch (Urbani et al., 1998). In spider crab *Inachus phalangium* the sperm of the last male is usually sufficient to fertilize more than one brood (Diesel, 1990). It is unknown, whether the sealed sperm of earlier males can be re-mobilized by the female if the sperm of the last male is exhausted (Pardo et al., 2013). Sperm displacement seems to be more effective in some brachyurans than in others but even in the most effective species sealing occasionally fails, resulting in multiple paternity in otherwise unipaternal species (Sainte-Marie et al., 2000; Jensen and Bentzen, 2012; Pardo et al., 2013; Rojas-Hernandez et al., 2014).

In long-lived species like clawed lobsters and some crabs, sperm can be stored across molts and utilized for years (Factor, 1995; Jensen and Bentzen, 2012; Pardo et al., 2013). This is made possible by sperm storage in mesodermal sectors of the seminal receptacles that are not shed during ecdysis (Becker et al., 2011; Pardo et al., 2013). For example, *Chionoecetes bairdi* females isolated from males after copulation produced viable eggs by 100% in the mating year and 97% and 71% in the following two years (Paul, 1984). Long term sperm storage by the females enables posthumous paternity, the ongoing siring of offspring after the death of males. In exploited species, in which males are selectively fished, posthumous paternity may even be vital for a sustainable fishery (Sainte-Marie et al., 2008; Pardo et al., 2013).

## FERTILIZATION TENT IN FRESHWATER CRAYFISH

Prior to spawning, freshwater crayfish produce a tent-like compartment on the underside of their body, which facilitates fertilization of the eggs and attachment of the zygotes to the pleopods. This structure is formed by a gelatinous secretion from the glair glands and persists for several hours. The glair glands develop in the weeks before spawning (Thomas and Crawley, 1975) and are thus good indicators of forthcoming egg-laying. They appear as creamy-white patches in the last thoracic sternal plates, the sterna and pleura of the pleon, the pleopods and the uropods (Fig. 4A). Glair glands are structurally different from other integumental glands and terminate with many pores on the underside of the thorax and pleon (Andrews, 1904; Thomas and Crawley, 1975).

**Fig. 4.**
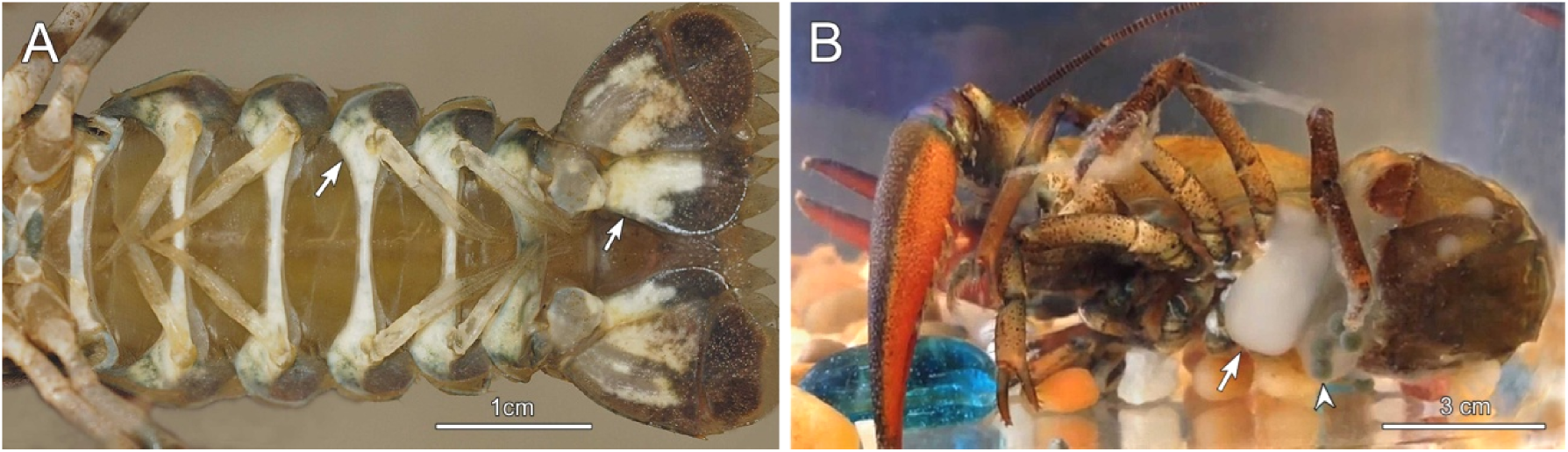
Glair glands and fertilization tent in crayfish females. (A) Whitish glair glands (arrows) on ventral side of pleon of *Orconectes limosus* (photo: Lucian Parvulescu). (B) Fertilization tent (arrow) in egg-laying *Pacifastacus leniusculus* composed of a gelatinous mass from the glair glands. A parchment-like sheet is formed at the contact zone with the water, which retains the eggs (arrowhead) within the tent (from a video by Bat Won).

Shortly before egg laying, crayfish form a ventral pouch by bending the pleon towards the underside of the cephalothorax. This pouch is then filled with the secretion from the glair glands (Fig. 4B), which in *Orconectes limosus* lasts about 30 minutes (Andrews, 1906). Lying on her back, the female then releases the eggs into the gelatinous mass and fertilizes them by sperm that is mobilized from the externally attached spermatophores or the annulus ventralis, depending on crayfish family (Mason, 1970; Gherardi, 2002). In *Orconectes limosus* egg laying takes about 10-30 min but in other species it can take hours (Andrews, 1904). The inner mass of the fertilization tent has a relatively low viscosity but at the contact zones with the water a more rigid parchment-like structure is formed, resembling the flysheet of a tent. Within this tent the eggs are transferred backwards by paddling movements of the pleopods and fastened to the oosetae of the pleopods. The attachment process requires repeated sideward turnings, which usually last for hours (Andrews, 1904, 1906; Mason, 1970).

The function of the fertilization tent may be manifold. It may protect the soft and highly labile fresh eggs from the osmotic stress of fresh water. Crayfish eggs achieve their typical round shape and rigid consistency only after hours when the outermost shell layer is hardened (Andrews 1904, 1906). Likewise, sperm are mostly unable to osmoregulate (Clark, 1978), and therefore, freshwater crustaceans with external fertilization may require a specific milieu for fertilization. The fertilization tent may also prevent egg loss during the complicated and long-lasting turnings that are necessary for egg attachment.

Glair glands and fertilization tents occur in all freshwater crayfish but are absent in marine lobsters, the closest relatives of crayfish (Factor, 1995). Therefore, they may be interpreted as a special adaptation to reproduction in fresh water. Freshwater crabs have internal fertilization and do not require such a protective mechanism but in freshwater shrimps and aeglids the eggs are externally fertilized as in crayfish (Chow et al., 1982; Almerão et al., 2010). However, there is no fertilization tent-like structure in these taxa objecting the above hypothesis. Hence, the fertilization tent has to be regarded as a unique feature of freshwater crayfish that has evolved in their stem group already.

## BROODING OF OFFSPRING IN SPECIALIZED BODY COMPARTMENTS

The simplest pattern of reproduction in Crustacea is the release of eggs and sperm into the water (broadcasting strategy) and larval development in the plankton. However, most crustacean groups have evolved mechanisms to brood their embryos, larvae or first juvenile stages (Gruner, 1993; Thiel, 2000, 2003). In some clades, for instance the Peracarida, brood care is obligatory and morphologically so specific that it can be used as a character for phylogenetic analysis (Richter and Scholtz, 2001) but in other lineages like the Euphausiacea brood care is limited to a fraction of the species only (Gómez-Gutiérrez, 2003). Here, I focus on brooding of the offspring in specialized body compartments of the mother. Information on brood care in dwellings that is based on behavioural interactions of mother and offspring rather than on structural specialities is found in reviews by Thiel (2003, 2007) and Vogt (2013).

The simplest type of brood care in crustaceans is carrying of the eggs on the mother until hatching of the nauplius larvae. Examples are the Copepoda among the Entomostraca and the Euphausiacea among the Malacostraca. Both groups include broadcasting and sac-spawning species. Species of the krill genera *Nyctiphanes* and *Nematoscelis* brood their eggs in a membranous sac on the thoracopods (Fig. 5A) (Gómez-Gutiérrez, 2003), whereas cyclopoid and some calanoid copepods carry the eggs in membranous sacs on the urosome (Fig. 10I) (Kiørboe and Sabatini, 1994).

**Fig. 5.**
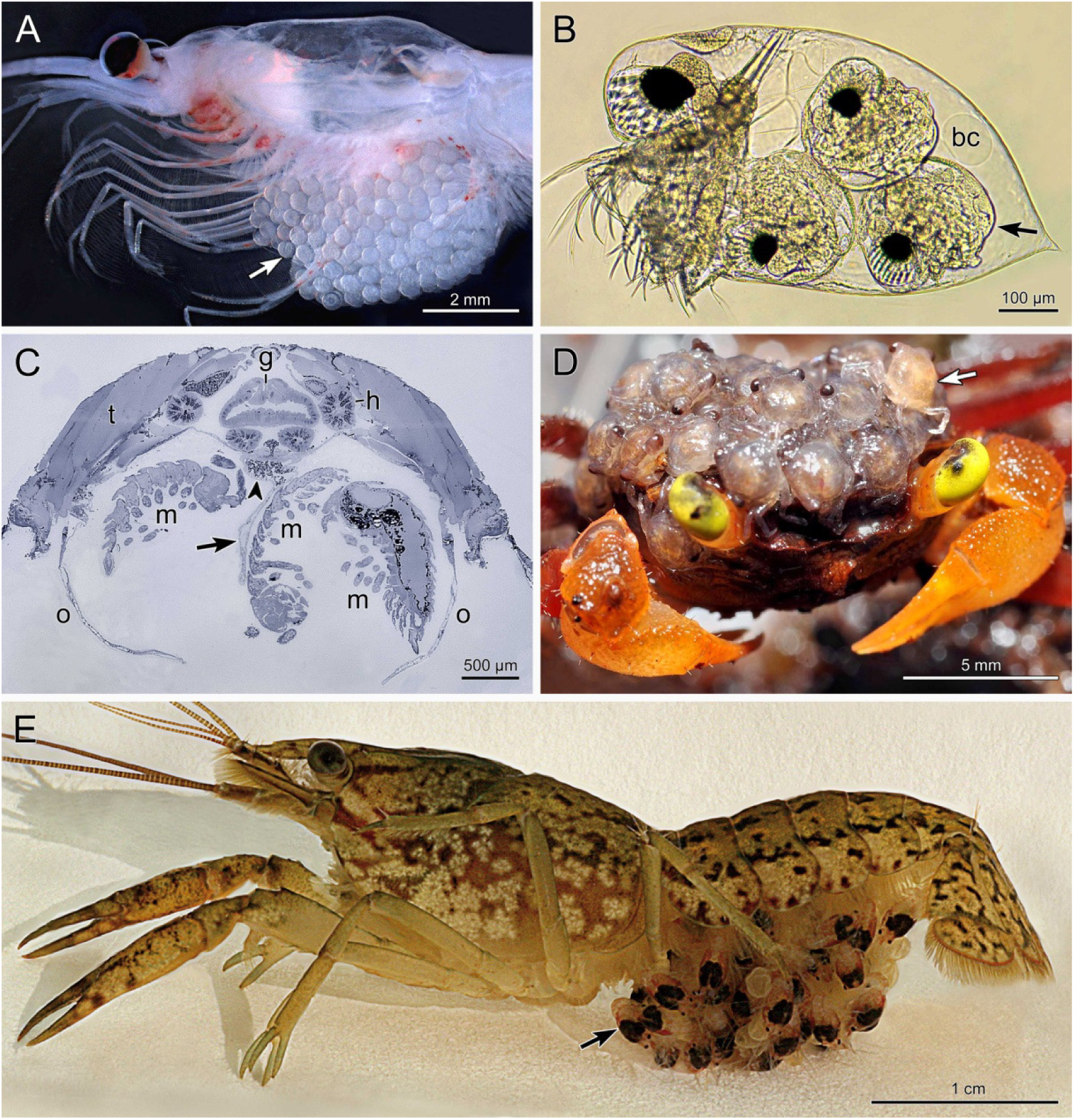
Brooding of eggs and posthatching stages in specialized body compartments of crustaceans from different environments. (A) Marine euphausiacean *Nyctiphanes australis* with egg sac (arrow) between thoracopods (photo: Anita Slotwinski). (B) Marine water flea *Evadne nordmanni* with advanced embryos (arrow) in huge dorsal brood chamber (bc) (photo: Maurice Loir). (C) Cross section through marsupium of terrestrial isopod *Cylisticus convexus* with brooded postlarval mancas (m). The marsupium is delimited by oostegites (o) and includes liquid and nutrient secreting cotyledons (arrow). Arrowhead denotes lipid droplets in upper part of cotyledon. g, gut; h, hepatopancreas; t, tergite (from Csonka et al., 2015). (D) Terrestrial crab *Geosesarma notophorum* carrying juveniles (arrow) on top of carapace (photo: Oliver Mengedoht). (E) Freshwater crayfish *Procambarus virginalis* carrying stage-2 juveniles (arrow) on pleopods (from Vogt and Tolley, 2004).

**Fig. 6.**
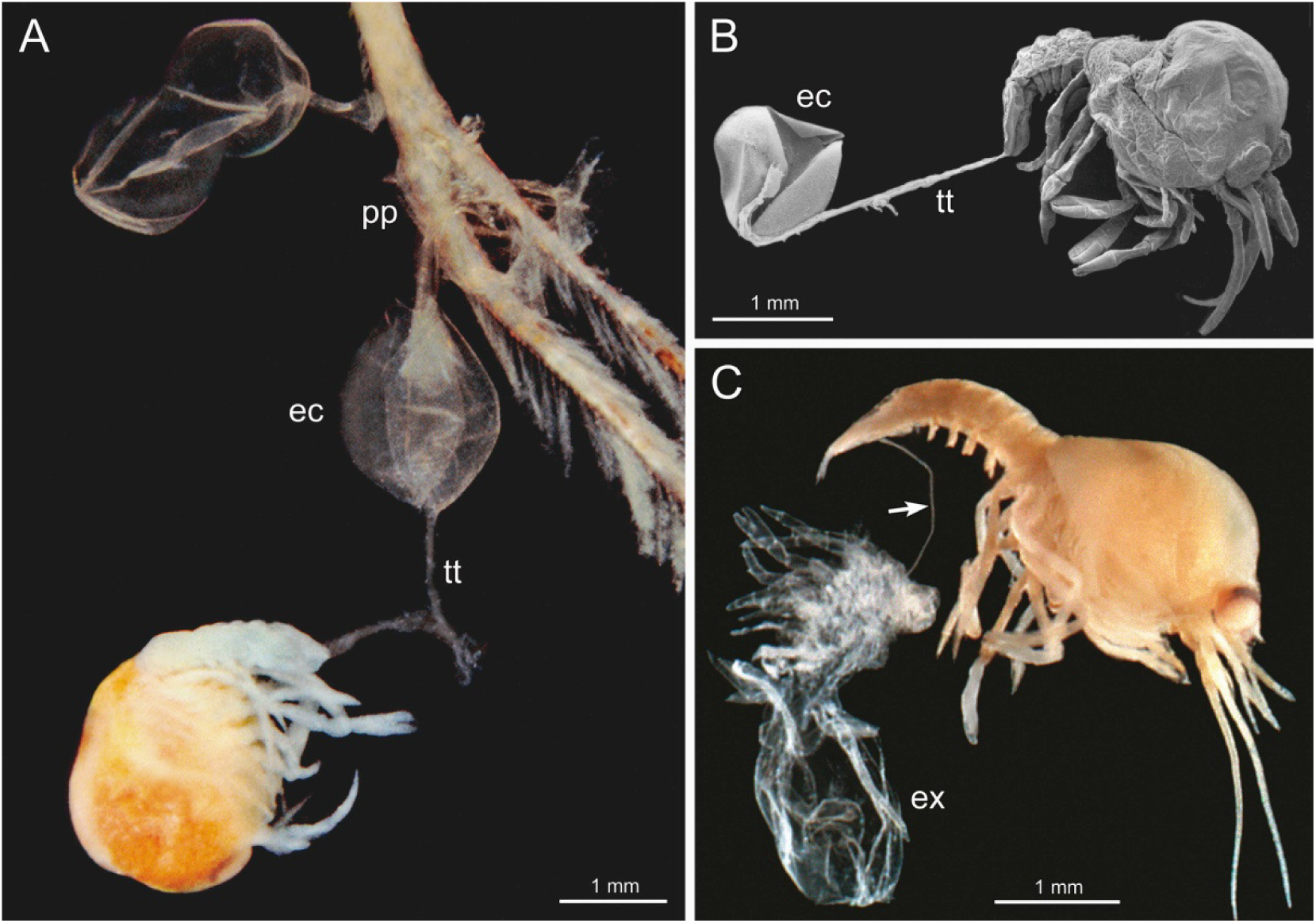
Safety lines in brooded crayfish juveniles. (A) Safeguarding of hatching by telson thread in *Procambarus virginalis*. The hatchling is secured to the maternal pleopod (pp) via telson thread (tt) and egg case (ec) (from Vogt and Tolley, 2004). (B) Scanning electron micrograph of telson thread connection between hatchling and egg case (from Vogt and Tolley, 2004). (C) Safeguarding of first molting by anal thread in *Procambarus virginalis*. The anal thread (arrow) secures the newly emerged stage-2 juvenile to its exuvia (ex), which *in situ* is hooked in pleopodal structures of the mother (from Vogt, 2008).

**Fig. 7.**
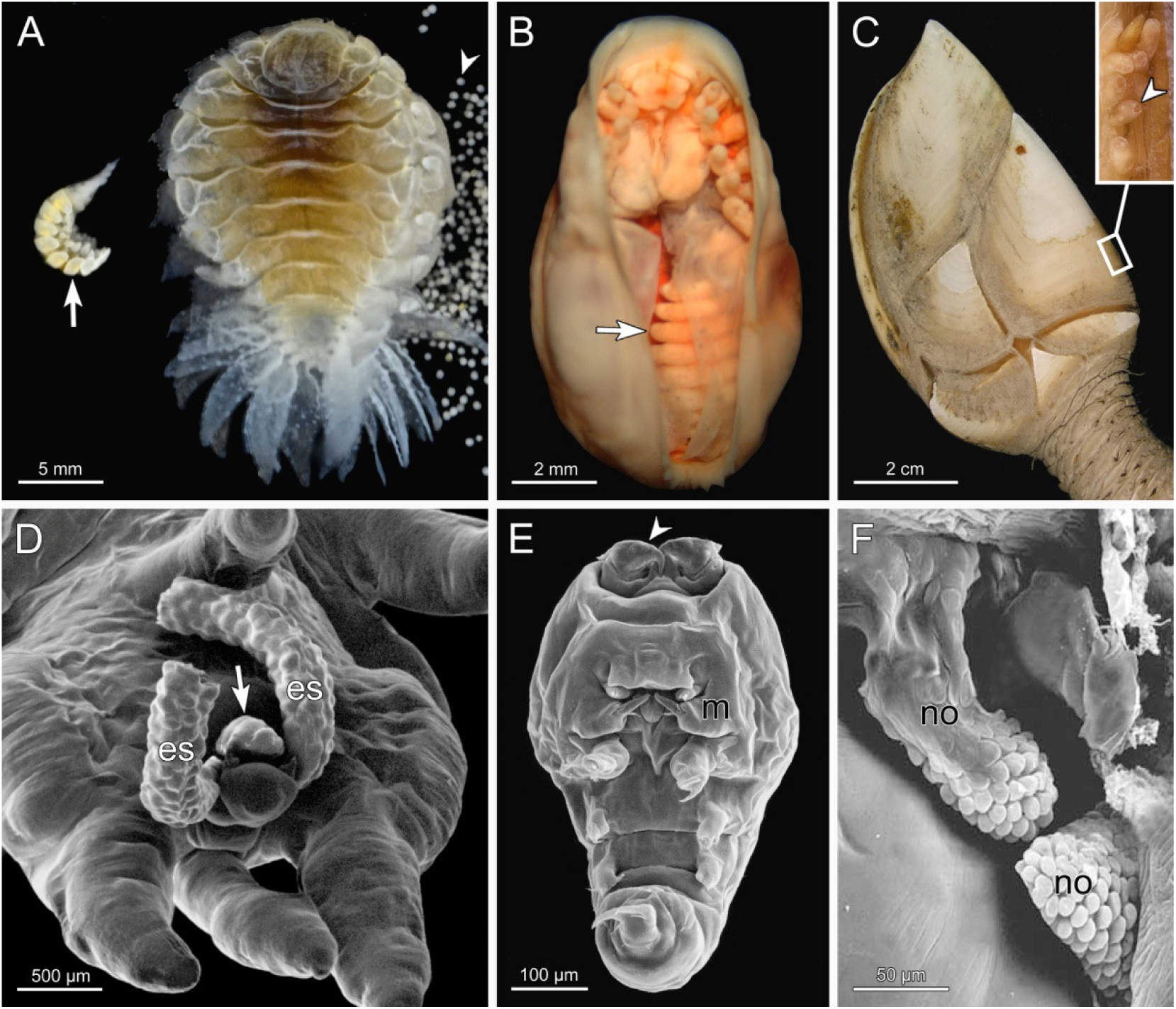
Dwarf males in parasitic Isopoda (A, B) and Copepoda (D-F) and sessile Cirripedia (C). (A) Dorsal view of ovigerous female and lateral view of male (arrow) of *Pseudione overstreeti*. Arrowhead denotes detached eggs (photo: Brent P. Thoma). (B) Ventral view of female *Zonophryxus quinquedens* with attached male (arrow) (from Raupach and Thatje, 2006). (C) Female gooseneck barnacle *Trianguloscalpellum regium* with several tiny males (arrowhead) in receptacle inside scutal edge (frame) (from Yusa et al., 2012). (D) Ventral aspect of posterior body part of female *Chondracanthus lophii* with tiny male (arrow) attached between egg sacs (es) (from Østergaard and Boxshall, 2004). (E) Close-up of male showing hooks on antennae (arrowhead). m, maxilla (from Østergaard, 2004). (F) Pinecone-like nuptial organs (no) of female serving as holdfast and nutrient source for the male (from Østergaard and Boxshall, 2004).

**Fig. 8.**
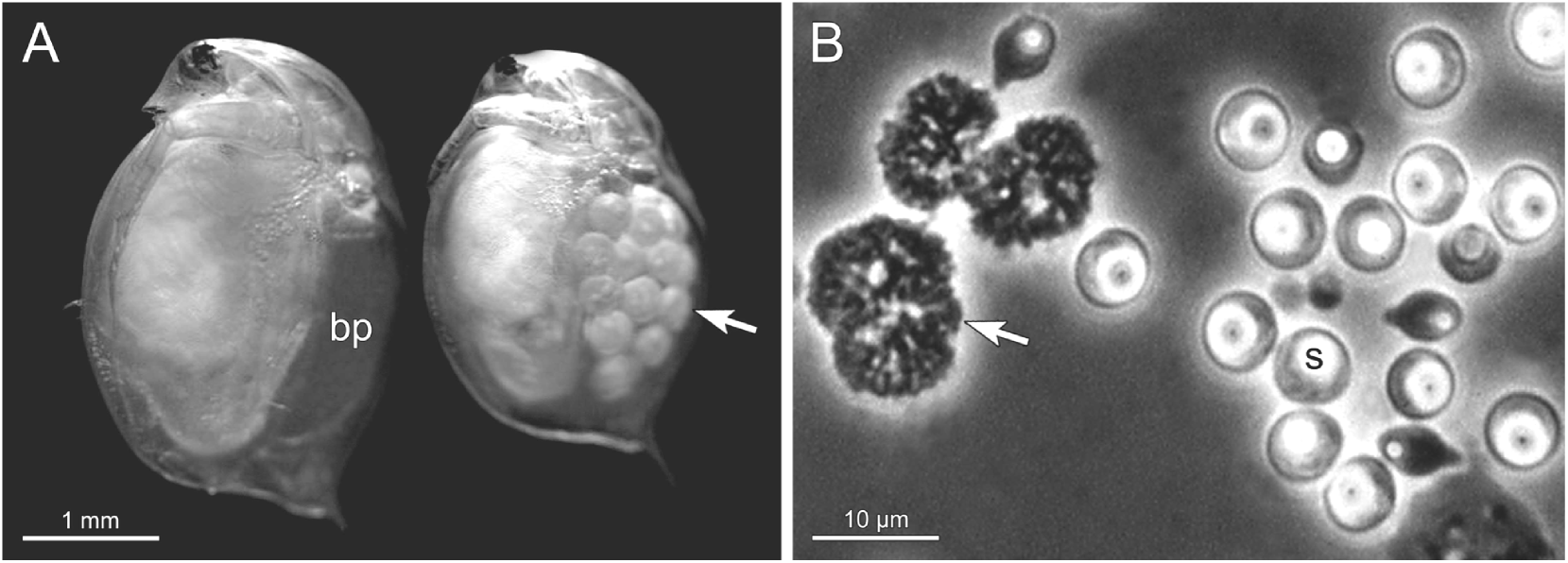
Bacteria-induced castration and gigantism in water flea. (A) Comparison of healthy (right) and *Pasteuria* ramosa-infected (left) *Daphnia magna*. The healthy specimen carries a clutch of eggs (arrow) in the brood pouch. The infected specimen is considerably larger and its brood pouch (bp) is empty indicating gigantism and castration (photo: Clay Cressler). (B) Cauliflower-type microcolony (arrow), grape-seed like stages and mature spores (s) of *Pasteuria ramosa* (photo: Dieter Ebert).

**Fig. 9.**
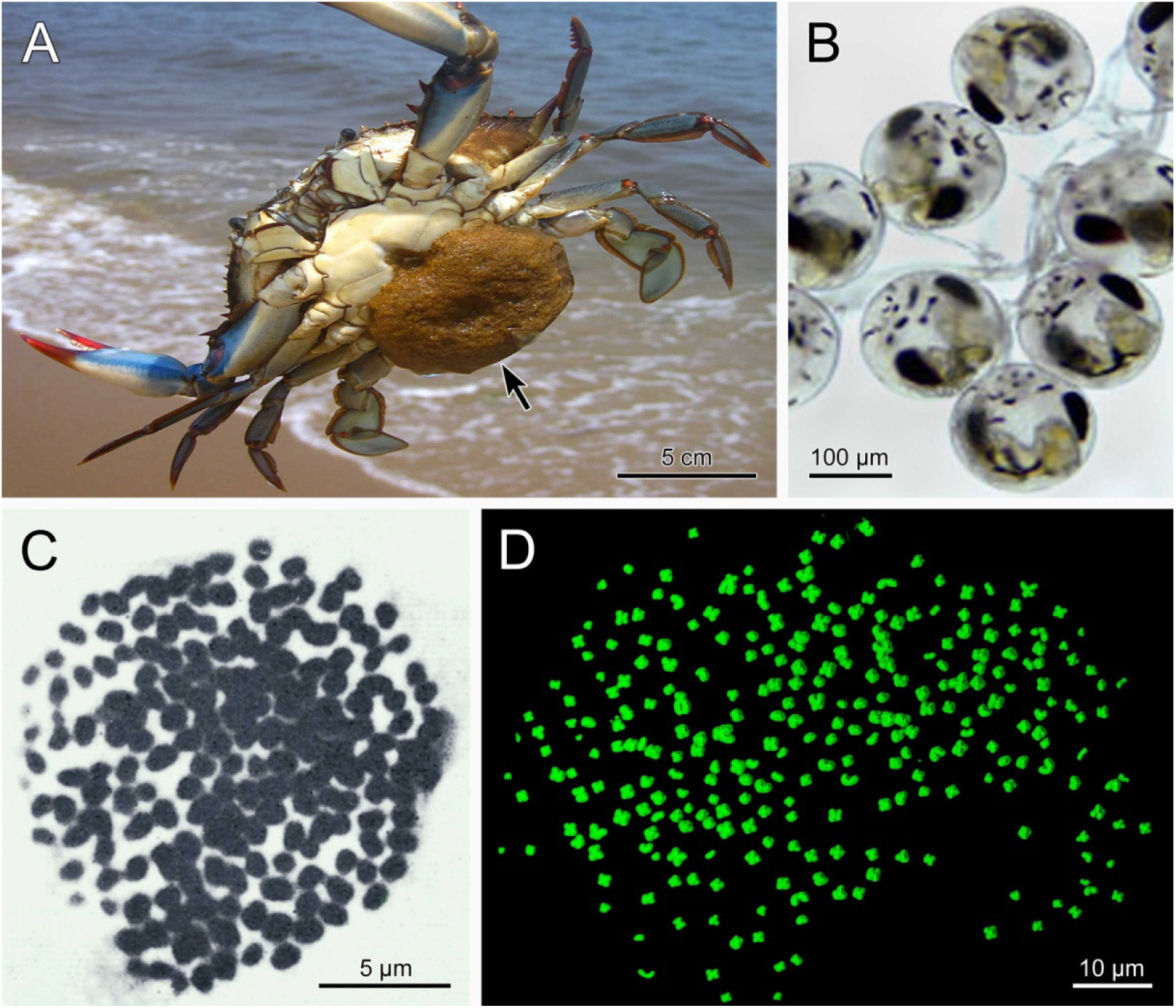
Record-breaking clutch size (A, B) and chromosome number (C, D) in Decapoda. (A) Blue crab *Callinectes sapidus* with "sponge" (arrow) including up to 8 million eggs (photo: Joe Reynolds). (B) Late embryos from sponge of blue crab (photo: Thomas H. Shafer). (C) Metaphase plate of primary spermatocyte of *Pacifastacus leniusculus trowbridgii* with 188 chromosomes (from Niiyama, 1962). (D) Metaphase spread of embryonic body cell of triploid crayfish *Procambarus virginalis* including 276 chromosomes (from Martin et al., 2016).

**Fig. 10.**
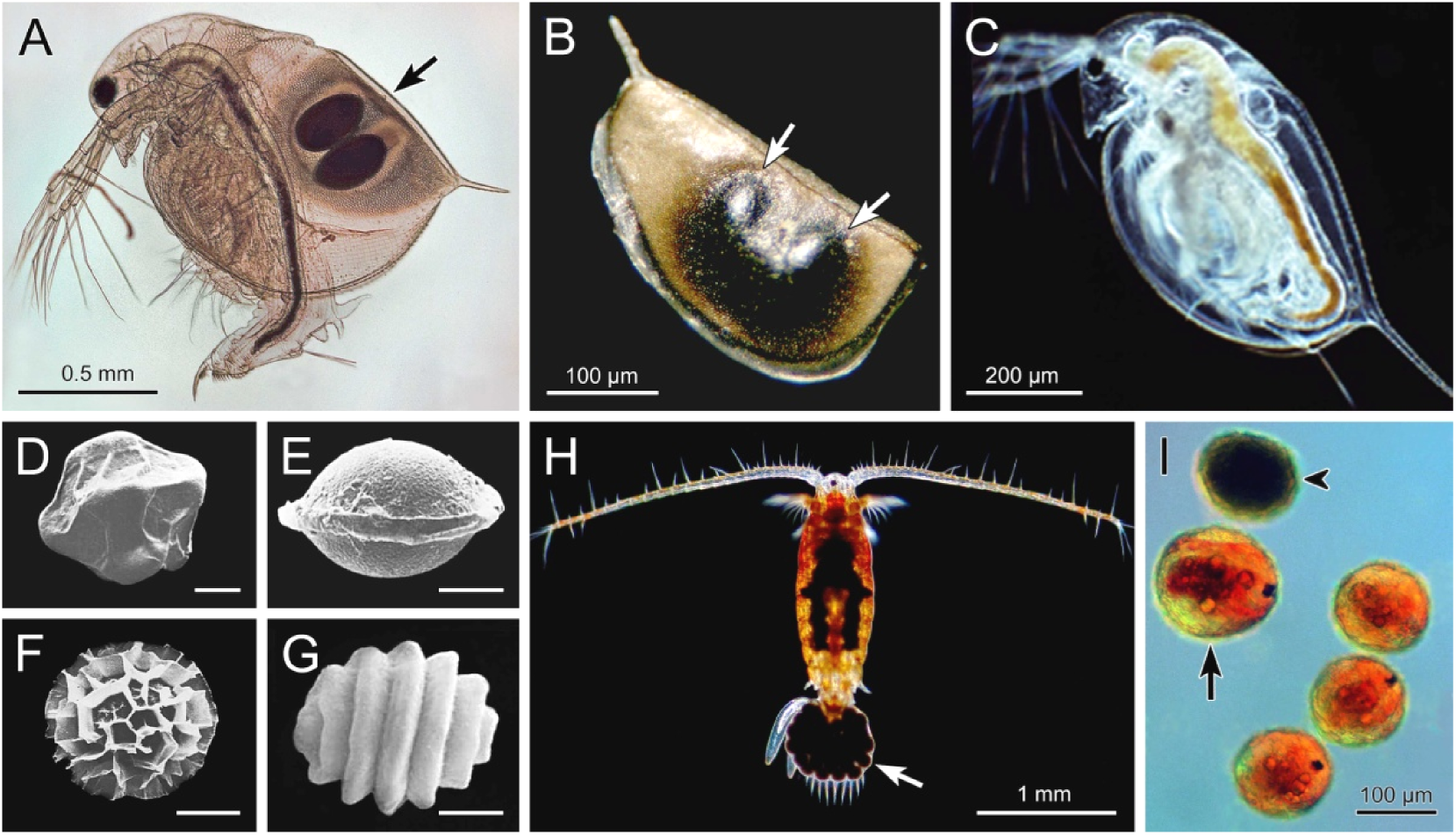
Record breaking viability of dormant stages in short-lived Crustacea. (A) Production of ephippium (arrow) in brood chamber of *Daphnia curvirostris* (photo: Jean-Pierre Claes). (B) Ephippium of *Daphnia pulicaria*, the branchiopod record holder of egg viability, including two eggs (arrows) (from http://www.upayan.info). (C) *Daphnia pulicaria* hatched from a ~700 year old dormant egg (photo: Dagmar Frisch). (D-G) Examples of specially shaped and ornamented dormant eggs of Branchiopoda. Bars: 50 μm (from Thiery and Gasc, 1991). (D) *Branchipus schaefferi*. (E) *Tanymastix stagnalis*. (F) *Chirocephalus diaphanus*. (G) *Imnadia yeyetta*. (H) *Onychodiaptomus sanguineus*, the copepod record holder of dormant egg viability with 332 years. Arrow denotes egg sac (photo: Ian Gardiner). (I) Diapausing eggs of *Onychodiaptomus sanguineus* from deeper lake sediment layer in blastula (arrowhead) and nauplius (arrow) stages (photo: Colleen Kearns).

Brooding of embryos and posthatching stages in dorsal brood chambers is typical of some entomostracan groups like the Cladocera (Fig. 5B) and Ostracoda (Fig. 11G). In cladocerans, the brood chamber is the space between the dorsal side of the trunk and the carapace. The eggs are laid in this pouch and the embryos develop until a miniature adult is released (Fig. 5B) (Mittmann et al., 2014). Some cladocerans with small eggs secrete a fluid rich in nutrients into this chamber, which may be absorbed by the eggs and developing embryos (Schminke, 2013). Interestingly, a dorsal brood pouch is also found in the malacostracan Thermosbaenacea, which raise the embryos and four posthatching stages in this chamber (Olesen et al., 2015). Usually, malacostracans brood their offspring on the ventral side of the body (Fig. 5C, E).

**Fig. 11.**
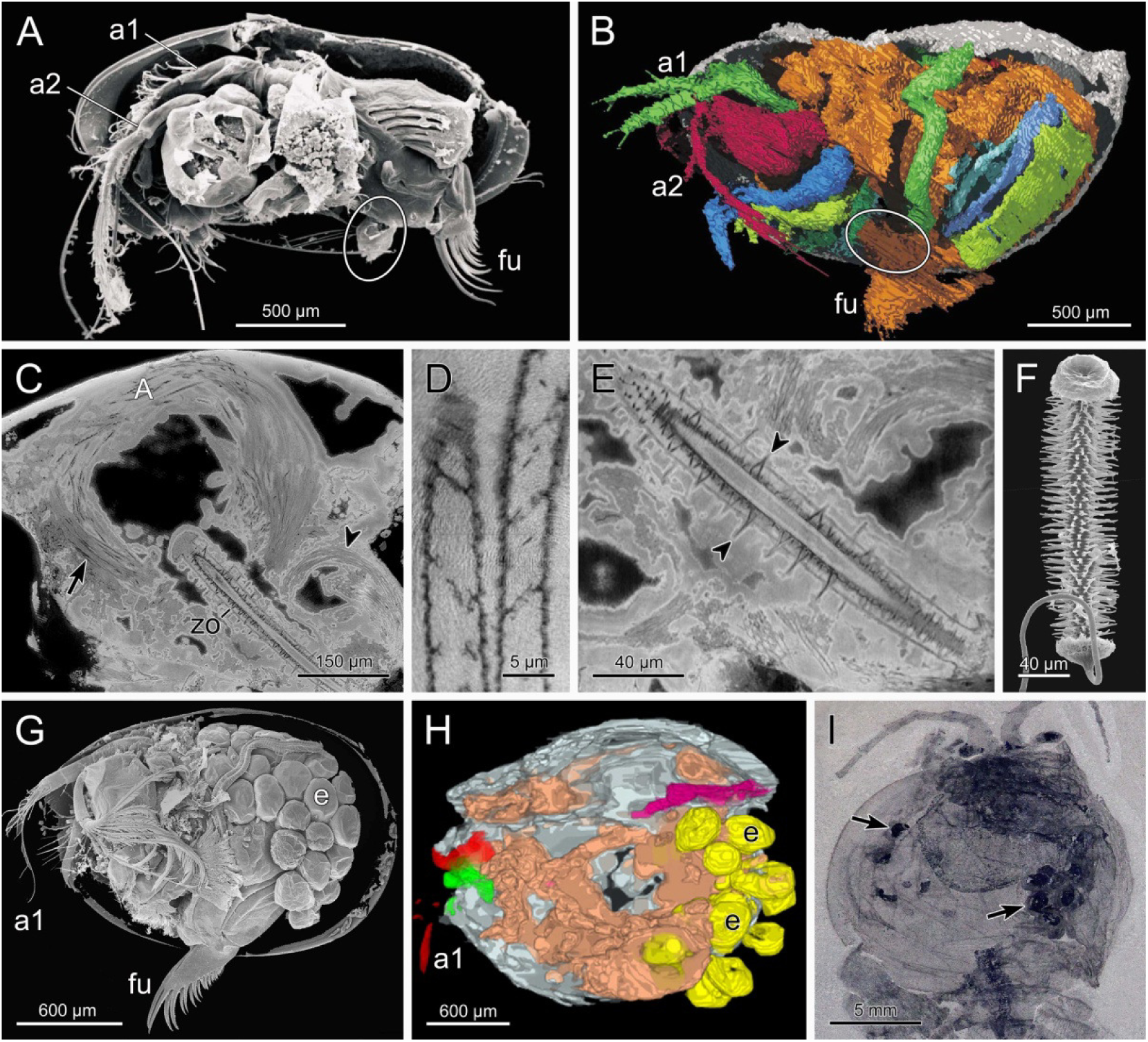
Record-breaking fossil ages of reproductive structures in Crustacea. (A, B) Penis (from Siveter et al., 2003). (A) Extant myodocopid *Xenoleberis yamadai* male showing ventrally located copulatory organ (encircled). al, first antenna; a2, second antenna; fu, furca. (B) Virtual reconstruction of 425 million-year-old myodocopid *Colymbosathon ecplecticos* male showing soft body parts including the copulatory organ (encircled). (C-F) Giant sperm and sperm pump (from Matzke-Karasz et al., 2014). (C) Lateral tomographic slice through 16 million-year-old *Heterocypris collaris* male showing bundles of giant sperm in seminal vesicle (arrow) and vas deferens (arrowhead). zo, Zenker organ acting as sperm pump. (D) Close-up of giant spermatozoa showing longitudinal spiralisation. (E) Close-up of Zenker organ with chitinous spines (arrowheads). (F) Zenker organ of extant *Heterocypris barbara* for comparison. (G-I) Brooded embryos. (G) Female of extant ostracod *Vargula hilgendorfii* showing embryos (e) in posterior brood chamber (from Siveter et al., 2014). (H) Oblique ventral view of 450 million-year-old ostracod *Luprisca incuba* with brooded embryos (arrows) (from Siveter et al., 2014). (I) Dorsal view of 508 million-year-old crustacean relative *Waptia fieldensis* showing lateral brood chambers with eggs (arrows) (from Caron and Vannier, 2016).

The Peracarida have a ventral brood chamber, the marsupium, which is delimited by specialized parts of the thoracopods, the oostegites (Fig. 5C). This marsupium probably evolved for mechanical protection of eggs and developing embryos in the sea but has become particularly important in terrestrial isopods, where it serves as a micro-aquarium for development (Hoese and Janssen, 1989; Csonka et al., 2015). It contains cotyledons that secrete the marsupial fluid and probably supply the young with nutrients. The latter possibility was inferred from the presence of lipid globules in the cotyledons (Fig. 5C) and the continuity of these organs with the lipid storing fat body of the mother (Hoese and Janssen, 1989).

In the Decapoda, the simplest type of brood care is carriage of the eggs until hatching. It is rare in Dendrobranchiata, the most basal decapods, but obligatory in all Pleocyemata. Dendrobranchiate shrimps are usually broadcast spawners and only the pelagic *Lucifer* species carry their eggs on the 3rd thoracopods until hatching of the nauplii (Lee et al., 1992). The pleocyemates brood their eggs on the pleopods at least until hatching of the zoea larva. Carriage of the embryos can last up to 16 months as shown for the American lobster *Homarus americanus* (Goudeau et al., 1987), and therefore, egg attachment must be very firm. This requirement is fulfilled by a special egg attachment system that consists of the rigid egg envelope, a firm but elastic egg stalk and the oosetae that arise laterally from the pleopods. This highly effective egg attachment system presumably originated in the common ancestor of the Pleocyemata some 430 million years ago (Porter et al., 2005).

In many Pleocyemata, carriage of the young is expanded to late larval stages or even to the juvenile stage (extended brood care). For example, in freshwater crayfish the juveniles are brooded on the pleopods (Fig. 5E) until the first feeding stage, which is juvenile stage 2 or 3, depending on family. However, if shelters are scarce the brooding period is considerably prolonged (Vogt, 2013). Other decapod examples of extended brood care are the freshwater crabs and aeglid anomurans, which carry the brooded stages in a ventral pouch that is formed by forward folding of the pleon under the cephalothorax. An exceptional case is the terrestrial crab *Geosesarma notophorum*, which carries the juveniles on top of the carapace covered by a film of water (Fig. 5D) (Ng and Tan, 1995).

The main advantage of brood care is the increased survival and growth of the young. Disadvantages are the higher energy expenditure of the caring mothers and the pronounced reduction of the number of eggs, particularly in species with extended brood care (Anger, 2001; Vogt, 2013). Therefore, brood care should be favoured only under specific circumstances. The Decapoda are particularly suitable to examine this issue in depth because they include non-brooders and brooders of different degrees and have independently evolved extended brood care in several lineages. Interestingly, brooding until the juvenile stage was convincingly demonstrated for less than 30 polar and coastal representatives of the ~12.000 marine species but is supposed to occur in about 70% of the ~3.000 species of freshwater decapods (Vogt, 2013). This strong imbalance suggests that extended brood care in decapods has evolved in response to heavily fluctuating abiotic and nutritional conditions (Anger, 2001; Vogt, 2013).

## SAFEGUARDING OF HATCHING AND FIRST MOLTING BY SAFETY LINES

Freshwater crayfish have evolved unique safety lines, the telson thread and anal thread, which secure the offspring during the immobile and helpless phases of hatching and first molting. At other times the brooded juveniles can actively hold on to their mother with specialized hooks on their peraeopods (Vogt and Tolley, 2004; Vogt, 2008).

The telson thread occurs in all crayfish families, the Astacidae, Cambaridae and Parastacidae (Andrews, 1907; Scholtz, 1995; Scholtz and Kawai, 2002; Vogt and Tolley, 2004). It emerges during hatching and probably originates from two sources, a secretion and the detaching inner layer of the egg case. The telson thread extends from the posterior end of the telson of the hatchling to its egg case, which persists on the pleopods after hatching (Fig. 6A, B). During eclosion it keeps the helpless hatchling passively secured to the mother (Fig. 6A) preventing it from being dislodged by the water current. Thereafter, it secures the hatchling during its attempts to actively hook into pleopodal structures of the mother (Vogt, 2008).

The anal thread secures the first molting and occurs only in the Cambaridae and Parastacidae. It is composed of cuticular material and originates from delayed molting of the hindgut (Scholtz, 1995; Rudolph and Rojas, 2003; Vogt, 2008). It keeps the emerging stage-2 juvenile passively linked to its exuvia (Fig. 6C), which itself remains hooked in pleopodal structures of the mother. In vitro tests revealed that this curious anal thread-exuvia-pleopod connection is firm enough to protect the freshly molted juveniles from being washed away (Vogt, 2008). After some hours, the anal thread is disconnected by flapping movements of the juveniles, which are now attached to the maternal pleopods with their peraeopodal hooks (Vogt, 2008). A special situation is found in the parastacid species *Astacopsis gouldi* and *Astacopsis franklinii* which secure three juvenile stages by anal threads (Hamr, 1992).

## DWARF MALES IN SESSILE AND PARASITIC CRUSTACEANS

Dimorphism between females and males is common in crustaceans and results from secondary sex characteristics related to sperm transfer and brood care. The larger sex is sometimes the female and sometimes the male. An extreme form of sexual dimorphism is the reduction of males to dwarf males, which has independently evolved in sessile cirripeds and parasitic isopods, copepods and cirripeds. Dwarf males are much smaller than corresponding females and are usually equipped with special structures to hold on the female. Their internal organs except of the gonads are reduced to different degrees, depending on taxon. The advantage of dwarf males is the permanent availability of males at a relatively low physiological cost under conditions of limited mating opportunities (Vollrath, 1998).

In Isopoda, dwarf males are typical of the ten families of the Bopyroidea and Cryptoniscoidea, which parasitize on other crustaceans from shallow waters to the deep sea (Williams and Boyko, 2012). The males reside on the females but their feeding biology is largely unknown. An example is the bopyrid *Pseudione overstreeti* (Fig. 7A) that infests the branchial chamber of the Mexican ghost shrimp *Callichirus islagrande* (Adkison and Heard, 1995). The greatest total length of the females of this parasite is 19.1 mm and the greatest width is 14.4 mm. The corresponding values in males are 4.9 mm and 2.3 mm, respectively. The male usually attaches to the ventral side of the female's pleon. In the dajid *Zonophryxus quinquedens*, which parasitizes on the carapace of the deep sea Antarctic shrimp *Nematocarcinus longirostris*, the females and males have maximum lengths of 20 mm and 4.5 mm, respectively (Brandt and Janssen, 1994). The dwarf males live on the underside of the females (Fig. 7B).

In the Cirripedia, dwarf males have independently evolved in the sessile Thoracica and the parasitic Rhizocephala (Yusa et al., 2012). An example is the pedunculate gooseneck barnacles, which either reproduce by simultaneous hermaphroditism, by hermaphrodites and dwarf males (androdioecy) or by pure females and dwarf males (dioecy) (Yusa et al., 2012). Androdioecy occurs in shallow-water species and dioecy occurs in the deep sea and in symbioses. In the androdioecious *Scalpellum scalpellum* two to five dwarf males are usually attached to a large hermaphrodite. In the dioecious deep sea species *Trianguloscalpellum regium* the dwarf males are very small and are located in groups in a specific receptacle inside the scutal edge (Fig. 7C). Scalpellid dwarf males are lecithotrophic and non-feeding. They are shorter lived than their carrying females or hermaphrodites, and thus, the mothers must repetitively acquire new males during their lifetime (Høeg et al., 2016). It is largely unknown how these dwarf males fertilize the females but males of *Verum brachiumcancri* were shown to possess a long penis that reaches the mantle cavity of the female (Buhl-Mortensen and Høeg, 2013).

The rhizocephalan cirripeds are endoparasites in marine Decapoda. The body of the female is composed of two parts, a root-like interna that penetrates the host organs and serves for the absorption of nutrients and a sac-like externa that serves for reproduction. In many species, the eggs in the externa are fertilized by the sperm of dwarf males. These invade the mantel cavity of the female as very small trichogons of only 200 μm in length, migrate to specific receptacles close to the ovaries, molt at the entrance of the receptacle and seal it with the shed cuticle. Thus, each of the two receptacles of a female is inhabited by a single trichogon only, which produces sperm and remains together with the female until the end of life (Hoeg, 1991). In some rhizocephalans, the male cyprid larvae inject spermatogonia into the mantle cavity, omitting the dwarf male stage (Hoeg, 1991, 1992).

In the Copepoda, dwarf males have evolved in several parasitic families. These dwarf males are many times smaller than their corresponding females (Østergaard and Boxshall, 2004). In the Chondracanthidae, a family parasitic on marine fishes, the males attach to immature females at the second copepodite stage, complete their development on the female and remain there until they die. A maximum of eight males on a single female has been recorded but in most members of the family adult females rarely have more than one male attached. An example is *Chondracanthus lophii*, in which tiny males adhere near the female genital apertures (Fig. 7D) to so-called nuptial organs, which are special structures for holdfast and probably nourishment. The males use the claws on the transformed antennae (Fig. 7E) for attachment. The pinecone-like nuptial organs (Fig. 7F) contain glandular tissue and are assumed to produce a secretion to nourish the males. Adult males have well developed mouthparts (Fig. 7E) and a functional oesophagus and midgut but no anus, suggesting that they feed on easily digestible food stuff like mucus (Østergaard, 2004; Østergaard and Boxshall, 2004).

## PARASITE-INDUCED CASTRATION, FEMINIZATION AND GIGANTISM

Parasitic crustaceans and bacteria can significantly change the reproductive biology of crustacean hosts by manipulating sex or causing infertility. Well known examples for the former are parasitic isopods and cirripeds (Williams and Boyko, 2012; Schminke, 2013). In the ghost shrimp *Callichirus islagrande* the reproductive activity is obviously suppressed when infected with the isopod *Pseudione overstreeti*. In all of the more than 100 host-parasite associations investigated, the gonads of the hosts were greatly reduced (Adkison and Heard, 1995). Moreover, since only female shrimp were found to be infested some female morphotypes may represent primary males that have changed sex after parasitic manipulation. Among the parasitic isopods some Bopyridae and Dajidae and all Entoniscidae and Cryptoniscoidea are assumed to be parasitic castrators of their hosts (Williams and Boyko, 2012).

A well-known bacterial manipulator of crustacean reproduction is the intracellular alpha-proteobacterium *Wolbachia pipientis*. Bouchon et al. (1998) detected this bacterium in 19 species from eight terrestrial isopod families, in one limnic isopod and in two estuarine isopods. Later, it was also found in a limnic amphipod and a marine gooseneck barnacle (Cordaux et al., 2012). *Wolbachia* was shown to induce feminization in isopods by converting genetic males into functional females (Juchault et al., 1992; Rigaud et al., 1997, 2001). Transinfection experiments established that the susceptibility or resistance to sex conversion depends much on the combination of host species and *Wolbachia* strain (Cordaux et al., 2004). *Wolbachia* is usually vertically transmitted via the egg cytoplasm but some horizontal transmission has obviously occurred as well as demonstrated by genetic analysis. Horizontal transmission may explain the occurrence of *Wolbachia* in crustacean groups outside of the Isopoda. The induction of parthenogenesis by *Wolbachia* as observed in insects (Stouthamer, 1997) has not yet been found in crustaceans.

A good example of castration and concomitant induction of gigantism in a crustacean host is the *Daphnia magna–Pasteuria ramosa* system (Ebert et al., 1996, 2004; Ebert 2005; Cressler et al., 2014). *Daphnia* females continue to produce eggs within the first days after infection by the bacterial manipulator (Ebert et al., 1996) but thereafter most of the infected individuals are castrated (Fig. 8A) and direct a considerable proportion of nutrients and energy towards reproduction of the parasite. At death, each host is filled with 10 to 20 million spores (Ebert, 2005). By manipulating food levels during the infection, Ebert et al. (2004) showed that both antagonists are resource-limited and that there is a negative correlation between host and parasite reproduction. Curiously, a certain proportion of the saved energy is channelled into growth of the host resulting in gigantism (Fig. 8A). Although illogical at first glance, the parasite benefits from this resource allocation because it can produce more spores in a bigger host (Ebert et al., 2004). *Pasteuria ramosa* has a polymorphic life cycle beginning with cauliflower-like rosettes and ending with individual spores (Fig. 8B). It is horizontally transmitted through spores that are released from dead host bodies into the water.

## RECORD-BREAKING CLUTCH SIZE AND CHROMOSOME NUMBER

Crustaceans are among the animals with the highest number of eggs and brooded embryos per clutch. Particularly large clutches are produced by big-sized brachyurans (Hines, 1991). The blue crab *Callinectes sapidus* carries in average about three million eggs in a sponge-like structure under the pleon (Fig. 9A, B). In large females this sponge was shown to contain up to 8 million eggs that are brooded until the zoea stage (Prager et al., 1990). In some *Cancer* species, the amount of eggs produced per female and lifetime exceeds 20 million (Hines, 1991). The blue crab, which has up to 18 broods (Graham et al., 2012), may even top this value.

The animal record holder in chromosome number is the freshwater crayfish *Pacifastacus leniusculus trowbridgii* (Niiyama, 1962). It has a diploid set of 376 chromosomes corresponding to 188 chromosomes in the gametes (Fig. 9C). The second highest chromosome number of 276 per body cell (3n) was recently found in the triploid crayfish *Procambarus virginalis* (Fig. 9D) (Martin et al., 2016). This species reproduces by apomictic parthenogenesis, i.e. without meiosis (Martin, 2015; Vogt et al., 2015). Therefore, the mature oocytes in the ovary should include 276 chromosomes as well, making *Procambarus virginalis* the new world record holder with respect to chromosome number of gametes.

The particularly high chromosome numbers in freshwater crayfish are explained by whole genome duplication events in their early evolution. This idea is based on the observation that some crayfish species have the double and four-fold chromosome numbers of others (Lécher et al., 1995; Martin et al., 2016). Decapods have haploid chromosome numbers ranging from 27 to 188 and a polyploid index of 41.7%, which is very high when compared to other animal groups (Otto and Whitton, 2000). Ancient polyploidization is mainly supposed for the Astacidea, Palinuridae and Paguroidea. An argument against the polyploidization hypothesis comes from the absence of a positive correlation of chromosome number and genome size. For example, *Astacus astacus* has fewer chromosomes (2n=176) than *Orconectes virilis* (2n=200) (Martin et al., 2016) but has a much larger genome size (19.64 pg versus 4.69 pg) (Jeffery, 2015), suggesting that high chromosome numbers may at least partly result from evolutionary chromosome fragmentation.

## RECORD-BREAKING VIABILITY OF DIAPAUSING EGGS

Short-lived members of ephemeral water bodies and limnic and coastal plankton often produce dormant eggs that can survive adverse environmental conditions for years (Radzikowski, 2013). This special adaptation to unfavorable and changing environments has apparently evolved in the earliest metazoans already (Cohen et al., 2009). In Crustacea, diapausing eggs and cysts are produced by the Branchiopoda, Copepoda and Ostracoda (Fig. 10A–I). Under laboratory conditions dormant eggs of the anostracan *Branchinecta packardi*, the notostracan *Triops longicaudatus* and the conchostracan *Caenestheriella gynecia* remained viable for a minimum of 16, 14 and 8 years, respectively (Radzikowski, 2013). Eggs from accurately dated lake sediments of the cladocerans *Daphnia galeata and Daphnia pulicaria* and the copepods *Boeckella poppei* and *Onychodiaptomus sanguineus* even hatched after an estimated 125, 700, 196 and 332 years, respectively (Hairston et al., 1995; Cáceres, 1998; Jiang et al., 2012; Frisch et al., 2014). The ~700 years of *Daphnia pulicaria* are the metazoan record of egg viability.

Crustacean dormant eggs can be the result of parthenogenetic or bisexual reproduction, depending on taxon (Radzikowski, 2013). Cladocerans usually produce subitaneous summer eggs by parthenogenesis and dormant winter eggs (Fig. 10A) by sexual reproduction. Dormant eggs are tolerant to environmental stress like drying, freezing, UV radiation and mechanical damage. For example, the cysts of the anostracan *Artemia franciscana*, a common model of dormancy, can survive exposure to temperatures of −271°C and +100°C (Radzikowski, 2013). This stress tolerance is achieved by protecting coverings that are often shaped and ornamented in a group or species-specific manner (Fig. 10D–G) (Thiery and Gasc, 1991; Fryer, 1996). These coverings also permit passage through the guts of birds facilitating dispersal and colonization of isolated water bodies (Radzikowski, 2013). In cladocerans, the dormant eggs are often additionally enveloped by a cuticular ephippium (Fig. 10A, B). Stress resistance is further provided by cryoprotectants like trehalose and glycerol and molecular chaperones like heat shock proteins (MacRae, 2010; Radzikowski, 2013).

The production of long-lived diapausing eggs in crustaceans is a bet-hedging strategy that constitutes an ecological and evolutionary reservoir. Mobilization of this reservoir can help the actual population to respond to environmental changes by enhancing the genetic variation and species richness (Hairston, 1996; Cáceres, 1998). In Oneida Lake, New York, diapausing eggs accumulate in the sediments to densities of 2.5 x 10^4^ eggs/m^2^ for *Daphnia galeata* and 8.0 x 10^4^ eggs/m^2^ for *Daphnia pulicaria* (Cáceres, 1998).). ^210^Pb dating of sediments and resurging experiments revealed that the two *Daphnia* populations have persisted in the lake for >200 years and that the eggs remain viable for >125 years. Annual emergence rates back to the water column ranged between 0 and 25 *Daphnia*/m^2^. Because annual variation in the size of the overwintering water-column population ranged between 0 and 2.5 individuals/L, the contribution of emergence to the development of the spring population was considerable in some years and negligible in others (Cáceres, 1998). In copepod species, the density of diapausing eggs can be as high as >10^6^/m^2^ and their annual mortality rate can be as low as 1% (Hairston et al., 1995; Hairston, 1996).

Egg banks of crustaceans are in many ways analogous to the seed banks of terrestrial plant species and are valuable tools for ecological, biogeographical and evolutionary research. They are useful to study the change of plankton communities over time (Ohtsuki et al., 2015) and to reconstruct invasion histories (Mergeay et al., 2006). Egg banks are also suitable to investigate the influence of man-made eutrophication on plankton. For example, increasing resistance to toxic cyanobacteria that drastically proliferated after eutrophication of Lake Constance, Germany, was established in *Daphnia galeata* from different sediment layers (Hairston et al., 1999). Phenotypic comparison of recent and resurrected ancient *Daphnia pulicaria* from South Center Lake, USA, revealed significant shifts in phosphorus utilization rates after the beginning of eutrophication (Frisch et al., 2014). Recent individuals showed steeper reaction norms with high growth under high phosphorus and low growth under low phosphorous, while genotypes from pre-industrialized agricultural eras showed flat reaction norms, yet higher growth efficiency under low phosphorous. Moreover, Brede et al. (2008) demonstrated that anthropogenic alterations of habitats are associated with long-lasting changes in communities and species via interspecific hybridization and introgression.

## RECORD-BREAKING FOSSIL AGE OF REPRODUCTIVE STRUCTURES

Ostracod crustaceans provide one of the most complete and consistent fossil records of any animal group including tens of thousands of fossil species dating back to the Cambrian (Harvey et al., 2012; Rodriguez-Lazaro and Ruiz-Muñoz, 2012). Usually, only their shells are preserved but in the last decade some outstanding fossils with soft body parts have been detected. They revealed, among other things, record-breaking ages of copulatory organs, sperm and brooded embryos.

Most ostracods are sexually reproducing and transfer sperm with complex paired copulatory organs called hemipenes (Fig. 11A) (Cohen and Morin, 1990; Martens, 1998). In an exceptionally well-preserved specimen of the myodocopid *Colymbosathon ecplecticos* from the Lower Silurian of Herefordshire, England, 3-D reconstruction revealed amazing details of the soft body including a copulatory organ (Siveter et al., 2003). This ostracod was preserved as a three-dimensional calcite infill in nodules hosted within volcanic ash. Its copulatory organ is relatively large and stout (Fig. 11B), projects anteriorly and has lobe-like distal flanks. With an age of 425 million years it is the oldest penis documented for any animal.

Ostracods of the suborder Cypridocopina are famed for having some of the longest sperm in the animal kingdom as discussed above. Matzke-Karasz et al. (2014) have recently discovered fossil giant sperm by X-ray synchrotron microtomography of 16 million-year-old *Heterocypris collaris* and *Newnhamia mckenziana* from Queensland, Australia. Giant sperm bundles were found in the seminal vesicles and vasa deferentia of a male (Fig. 11C) and the sperm receptacles of females. These bundles included spermatozoa of excellent threedimensional preservation showing longitudinal coiling (Fig. 11D) and spiraling of the sperm nucleus. Other well-preserved reproductive structures in the male were the paired Zenker organs (Fig. 11E, F), which are chitinous and muscular pumps that help to transfer the sperm into the female (Yamada and Matzke-Karasz, 2012).

The sperm of *Heterocypris collaris* and *Newnhamia mckenziana* are the oldest giant sperm on record and the third oldest sperm of any animal. Older fossil sperm were found in a 50 million years old annelid cocoon from Antarctica (Bomfleur et al., 2015) and a spring tail trapped 40 million years ago in Baltic amber (Poinar, 2000). The giant sperm of ostracods are exceptional in as far as they are considered to have originated only once some 100 million years ago and have been retained since then, which stands in contrast to the generally rapid evolution of sperm (Smith et al., 2016). Indirect evidence of their occurrence already in the Cretaceous comes from the detection of Zenker organs, which are restricted to taxa with giant sperm (Matzke et al., 2009). The long history and persistence of giant sperm in ostracods makes them a unique model to study the evolutionary significance and function of this unusual sperm type in animals (Matzke-Karasz et al., 2014; Smith et al., 2016).

Direct evidence of brood care is rare in invertebrate fossils (Wang et al., 2015). In 2007, Siveter and colleagues detected a 425 million years old myodocopid ostracod, *Nymphatelina gravida*, in Herefordshire, England, which included 20 ovoid and two valve-shaped globules in the posterior domiciliar area resembling brood care in extant myodocopids (Fig. 11G). Accordingly, these 550 μm long globules were interpreted as brooded eggs and juveniles, which is a unique combination in fossil invertebrates (Siveter et al., 2007). Later, Siveter et al. (2014) found a pyritized ostracod, *Luprisca incuba*, with well preserved embryos in the Upper Ordovician of central New York State, USA (Fig. 11H), providing conclusive evidence of a conserved brood care strategy within the Myodocopida for at least 450 million years.

The oldest crustacean-like fossils showing evidence of brood care are the ~508 million-year-old shrimp-like *Waptia fieldensis* from the middle Cambrian Burgess Shale formation in Canada and the ~515 million-year-old bradoriid *Kunmingella douvillei* from the lower Cambrian Chengjiang Lagerstätte in China (Shu et al., 1999; Caron and Vannier, 2016). The latter fossil includes clusters of rounded bodies of 150 μm in a ventral chamber delimited by distal setae of the postantennular appendages. These globules are probably brooded eggs resembling brood care in the phyllocarid *Nebalia bipes* (Shu et al., 1999). *Waptia fieldensis* carried the eggs in two anterior chambers located dorsolaterally between the body and the inner surface of the carapace (Fig. 11I). Differences in size and elemental composition of the globules suggest the presence of eggs and further developed embryos that have already exhausted some of the yolk, which is unique in animal fossils. The relatively small clutch size of less than 25 eggs and the relatively large egg size up to 2 mm in *Waptia* contrast with the high number of small eggs found in *Kunmingella*, indicating diversification of brooding strategies soon after the Cambrian emergence of arthropods (Caron and Vannier, 2016).

## CONCLUSIONS

The various crustacean groups have evolved quite different structural solutions for similar reproductive requirements, which may provide valuable study material for comparative morphologists, students of bionics and evolutionary biologists. For example, cirripeds were able to bring into line conflicting sessility and direct sperm transfer by the evolution of a highly stretchable hydraulic penis, whereas the vagile decapods remained with solid cuticular copulatory organs. Another example is sperm, which occur in crustaceans in almost all possible variants from flagellate to aflagellate. The latter have compensated the lack of flagellae by sperm deposition at favorable sites and the invention of new mechanisms of sperm motility that allow penetration of the egg envelope. There are also different modes of paternity assurance, which contribute to the more general topic of sperm competition. Brooding of the offspring on the body has repeatedly evolved but different body compartments have become specialized to carry the young. Dwarf males are an example of convergent adaptation of parasitic and sessile taxa to limited mating opportunities. Some specialities have evolved only once in the Crustacea such as the fertilization tent and the safety lines in freshwater crayfish. The interesting question remains, why these features stayed rare despite their obvious advantages.

The Crustacea hold animal records with respect to relative penis length, sperm body length, clutch size, chromosome number, dormant egg viability, and fossil ages of penis, giant sperm and brood care. Such data may please the reader and find their way in the Guinness Book of Records but they may also contribute to the solution of puzzling evolutionary issues. For example, brood care has obviously evolved and diversified since the early radiation of the Crustacea but has become obligatory only in some crustacean groups. Giant sperm have evolved more than 100 million years ago but despite the high energetic cost they persisted until presently. The extremely high chromosome number in freshwater crayfish may point at chromosome multiplication and fragmentation as a pathway to speciation and the evolution of higher taxa.

The reproductive specialities and curiosities of crustaceans may also be of interest for ecologists and applied biologists. An example is long-term sperm storage and posthumous paternity in long-lived decapods, which may allow sustainable male-based fishery without losing genetic diversity of the exploited populations. Another application is the use of dormant eggs and resurged individuals of short-lived crustaceans to reconstruct pristine ecosystems prior to man-made pollution and to study the speed and extent of genetic and physiological adaptations to environmental changes.

## ACKNOWLEDGEMENTS

The author is grateful to the following colleagues for providing photographs and information: Carola Becker (Belfast, UK), Peter G. Beninger (Nantes, France), Rudolf Diesel (Seefeld, Germany), Dieter Ebert (Basel, Switzerland), Dagmar Frisch (Kingston, USA), Paolo Galeotti (Pavia, Italy), Ian Gardiner (Calgary, Canada), Nelson G. Hairston Jr. (Ithaca, USA), Jens Høeg (Copenhagen, Denmark), Erzsébet Hornung (Budapest, Hungary), Colleen M. Kearns (Ithaca, USA), Maurice Loir (Orange, France), Peer Martin (Berlin, Germany), Renate Matzke-Karasz (Munich, Germany), Oliver Mengedoht (Recklinghausen, Germany), William A. Nelson (Kingston, USA), Pia Østergaard (London, United Kingdom), Luis Miguel Pardo (Valdivia, Chile), Lucian Pârvulescu (Timişoara, Romania), Tiffany Penland (Raleigh, USA), Michael J. Raupach (Wilhelmshaven, Germany), Stefan Siebert (Davis, USA), David J. Siveter (Leicester, United Kingdom), Anita Slotwinski (Brisbane, Australia), Piet Spaak (Dübendorf, Switzerland), Brent P. Thoma (Lafayette, USA) and Jean Vannier (Villeurbanne, France).

